# No evidence for genome editing in mouse zygotes and HEK293T human cell line using the DNA-guided *Natronobacterium gregoryi* Argonaute (NgAgo)

**DOI:** 10.1101/091678

**Authors:** Nay Chi Khin, Jenna L. Lowe, Lora M. Jensen, Gaetan Burgio

## Abstract

A recently published research article reported that the extreme halophile archaebacterium *Natronobacterium gregoryi* Argonaute enzyme (NgAgo) could cleave the cellular DNA under physiological temperature conditions in cell line and be implemented as an alternative to CRISPR/Cas9 genome editing technology. We assessed this claim in mouse zygotes for four loci (*Sptb*, *Tet-1*, *Tet-2* and *Tet-3*) and in the human HEK293T cell line for the EMX1 locus. Over 100 zygotes were microinjected with nls-NgAgo-GK plasmid provided from Addgene and various concentrations of 5’- phosphorylated guide DNA (gDNA) from 2.5 ng/μl to 50 ng/μl and cultured to blastocyst stage of development. The presence of indels was verified using T7 endonuclease 1 assay (T7E1) and Sanger sequencing. We reported no evidence of successful editing of the mouse genome. We then assessed the lack of editing efficiency in HEK293T cell line for the EMX1 endogenous locus by monitoring the NgAgo protein expression level and the editing efficiency by T7E1 assay and Sanger sequencing. We reported that the NgAgo protein was expressed from 8 hours to a maximum expression at 48 hours post-transfection, confirming the efficient delivery of the plasmid and the gDNA but no evidence of successful editing of EMX1 target in all transfected samples. Together our findings indicate that we failed to edit using NgAgo.

## Introduction

Type II CRISPR/Cas9 genome editing system offers the ability to efficiently and precisely edit DNA using a combination of the Cas9 endonuclease enzyme and a single guide RNA gRNA [1]. However the requirement of a specific protospacer adjacent motif (PAM) sequence limits the ability of the Cas9 enzyme to edit any nucleotide of a genome. Recently a report described a novel genome editing technology based on the archaebacterium *Natrnobacterium gregoryi* Argonaute (NgAgo) enzyme. Gao et al. described an endonuclease activity for NgAgo, which has the ability to create a site-specific double strand break in the DNA under the guidance of the 24 nucleotide-5’-phosphorylated single stranded DNA (gDNA) which binds to the endogenous DNA [2]. Gao et al. demonstrated the high efficiency of NgAgo to edit the genome under physiological temperature condition without the requirement of a PAM [2]. Interestingly, Gao et al. demonstrated the intolerance of NgAgo to guide-target mismatches leading to negligible off-target effects. With such ability to target any nucleotide in the genome and with an equivalent efficiency to CRISPR/Cas9 genome editing technology, NgAgo has an undeniably important therapeutic potential [3]. We sought to assess the efficiency of NgAgo in two different systems: mouse zygotes and the HEK293T human cell line to determine the suitability of NgAgo as an alternative to the CRISPR/Cas9 genome editing tool. Here we report our attempts to edit the mouse and human genomes using NgAgo. We synthetized the gDNA and co-microinjected in mouse zygotes with the nls-NgAgo-GK plasmid vector provided by Gao et al. We also co-transfected the gDNA with a modified Flag-nls-NgAgo-GK plasmid into HEK293T cells and assessed the DNA editing using T7 endonuclease assay and Sanger sequencing. We monitored the expression of the protein for the first 48 hours post-transfection in HEK293T cells. We found no evidence for a double strand break and editing of the DNA under various conditions and optimizations. We concluded that we failed to edit the genome using NgAgo.

## Results and discussion

To assess NgAgo efficiency to create a double strand break under the guidance of a single gDNA as described in Gao et al. [2] we used mouse zygotes as a system model. We firstly co-injected mouse zygotes with the nls-NgAgo-GK plasmid and the gDNA targeting four different genes (*Sptb*, *Tet1*, *Tet2* and *Tet3*). The gDNA selected were previously shown to edit efficiently (over 60% efficiency) as an sgRNA with CRISPR/Cas9 mediated genome editing [4]. Initially, we targeted exon 26 of *Sptb* in mouse zygotes (Figure 1A). We titrated the gDNA at various concentrations (2.5, 25 or 50 ng/μl) and co-injected with 5, 10 or 15 ng/μl of nls-NgAgo-GK plasmid, provided by Gao et al. and available at Addgene, into the pronucleus of the fertilized zygotes. The zygotes were cultured for 4 days to the blastocyst stage of development. No abnormality in the development was found in these embryos. From the 49 bastocysts that were genotyped (Table 1), we found the amplification of a band corresponding to the expected amplicon length for *Sptb* (Figure 1B presented on NgAgo 5 ng/μl and 50 ng/μl of gDNA). We performed a T7 endonuclease assay (T7E1) and Sanger sequencing to identify indels. We could not identify any indels from the T7E1 assay (Figure 1C) and the Sanger sequencing (Figure 1D) on all blastocysts. We then hypothesized the lack of editing could be gene specific, hence we decided to assess three other genes; exon 5 of *Tet-1* (Suppl Figure 1A), exon 3 of *Tet-2* (Suppl Figure 1B) and exon 5 of *Tet-3* (Suppl Figure 1C) as described previously [4]. The zygotes were co-injected with 5ng/μl of nls-NgAgo-GK plasmid and 2.5 ng/μl of gDNA. We constantly found one band using gel electrophoresis corresponding to the expected amplicon size for *Tet-1* (10 blastocysts), *Tet-2* (14 blastocysts) and *Tet-3* (13 blastocysts). We performed a T7E1 assay and genotyped the blastocysts by Sanger sequencing. We found again, no evidence for developmental phenotype and presence of indels (data not shown). We further assessed the NgAgo editing efficiency for *Sptb* and *Tet-2* on pups born from the microinjection sessions. All pups displayed a normal phenotype and developed normally to adulthood. We performed a T7E1 assay and Sanger sequencing to assess the editing efficiency and we did not find any indels, suggesting no successful editing for *Sptb* and for *Tet-2* (Table 2). Together, this suggests that we failed to generate indels for the four analyzed mouse loci (*Sptb*, *Tet-1*, *Tet-2* and *Tet-3*) in over 86 mouse blastocysts and 25 mouse pups using NgAgo, suggesting NgAgo does not create a double strand break nor that it is capable of editing the mouse genome. We therefore speculated this lack of editing activity could be due to the degradation of the protein within the cell. To address this hypothesis, we tagged the protein with a flag tag upstream of the nls signal (Flag-nls-NgAgo-GK, Figure 1E) and monitored the expression of the protein and whether NgAgo edited the DNA in the HEK293T cell line at various time points from 8 to 48 hours post lipofection with or without the gDNA for exon 3 of the EMX1 gene (Figure 1F). We used an anti-flag antibody to probe for NgAgo expression, and utilized GAPDH as a loading control. We first conducted a PCR and a T7E1 assay on the NgAgo-EMX1-lipofectamine treated cells to determine whether the DNA was edited. We found no evidence for editing in any NgAgo-lipofection samples at 8 and 12 hours post-transfection (Figure 1G). Similarly, there was no editing for the additional time points of 24 and 48 hours post transfection (Supplementary Figure 2 A and B). We therefore speculated that the NgAgo protein could be not expressed post transfection or rapidly degraded after transfection and required a specific timing to edit the DNA. To verify this hypothesis, we followed the kinetics of the NgAgo protein production and degradation from 8 to 48 hours post transfection. We noticed the NgAgo protein expression started at 8 hours and persisted over 48 hours post transfection (Figure 1H), with maximum expression being observed at 48 hours post transfection compared to 8 hours (p = 0.02) (Figure 1H and Supplementary Figure 3) suggesting an efficient delivery of NgAgo and the gDNA. Interestingly, we noticed the presence of small-Flag-tagged fragments by 8 hours post transfection, suggesting the protein started degrading, with the fragment intensity reaching its peak at 48 hours post-transfection associated with the peak protein expression mentioned above with and without the co-transfection of the gDNA (Figure 1H). We noted no difference in protein expression and degradation with or without the gDNA (p = 0.0631 and p = 0.25 respectively) (Figure 1H and Supplementary Figure 3). Therefore, since degradation of the protein is the same between NgAgo treated samples with and without gDNA, the difference in protein expression is unlikely to be due to this rapid degradation.

**Figure 1:**
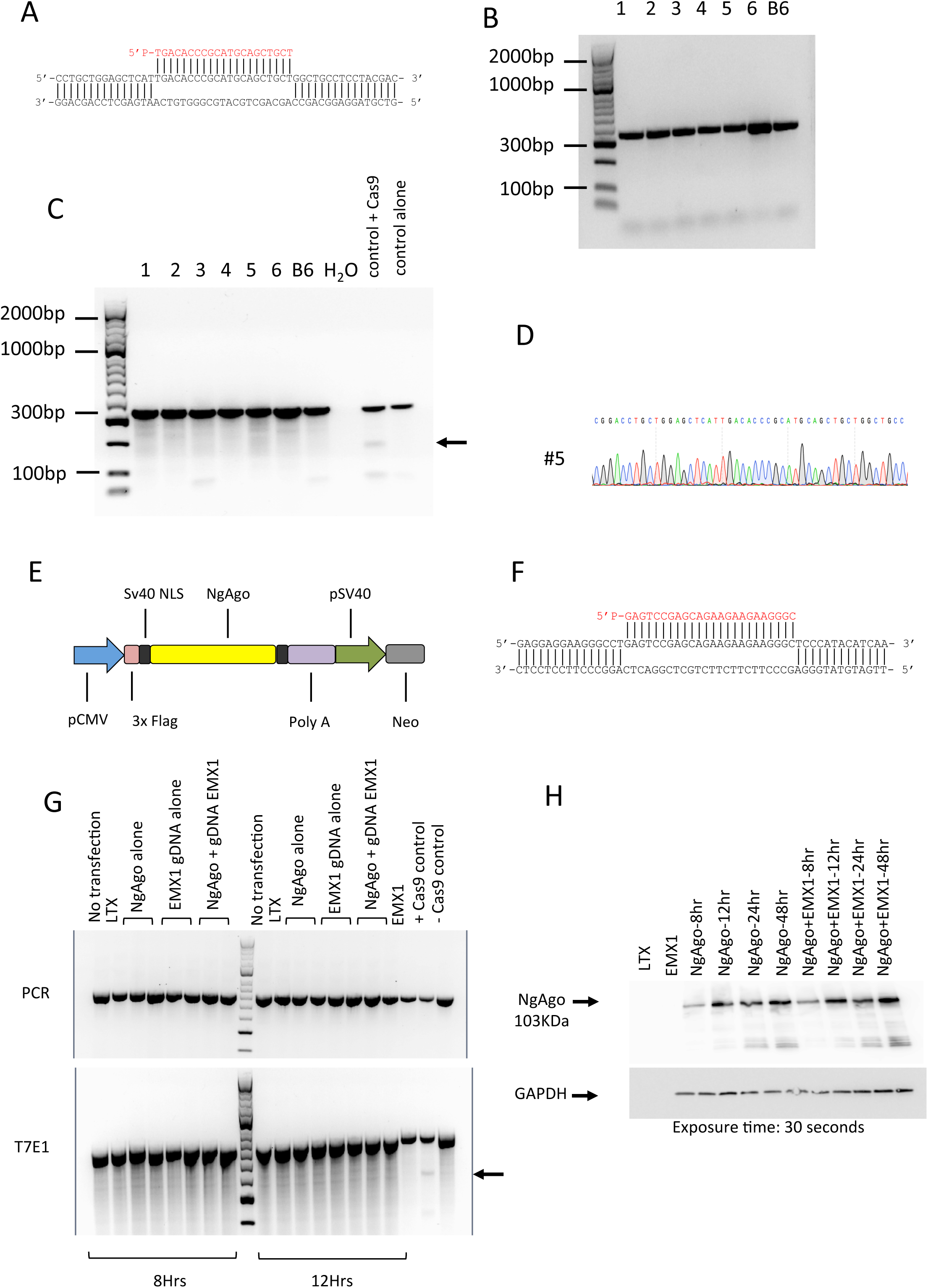
No evidence for double strand break cleavage and editing from NgAgo. A) DNA sequence indicating the locus targeted for the exon 26 of *Sptb*. The gDNA sequence is indicated in red. B) Gel electrophoresis (1.5%) of *Sptb* blastocysts (n=6) co-injected with nls-NgAgo-GK plasmid (2.5 ng/μl) and 2.5 ng/μl of gDNA. C57BL/6 DNA (B6) was also amplified as a control. The PCR product is 326 bp. C) T7 endonuclease 1 (T7E1) assay on the *Sptb* blastocysts indicating the absence of heteroduplexes suggesting indels in 6 blastocysts co-injected with nls-NgAgo-GK and gDNA. C57BL/6 (B6) non-edited control DNA was utilized as a negative control. A positive control in mouse zygotes edited with CRISPR/Cas9 was used as a positive control. The arrows indicate the presence of heteroduplexes suggesting a successful editing of the DNA. D) Representative chromatogram of one *Sptb* blastocyst (#5) suggesting no editing of the DNA under the DNA-guided NgAgo. E) Schematic diagram representing the flag-nls-NgAgo-GK plasmid. The expression of NgAgo is driven from a CMV promoter. A Flag tag was inserted in the 5’ end of NgAgo sequence. Two Sv40 nuclear localization signals were inserted in the 3’ end of the NgAgo sequence and in the 3’ end of the flag tag. A Ploy A tail was appended to the sequence in the 3’ end a Neomycin cassette was added in the 3’ end of the plasmid sequence. F) DNA sequence indicating the targeting of the exon 3 from EMX1 human sequence. The gDNA sequence is indicated in red. G) Gel electrophoresis (2%) of the PCR for EMX1 in HEK293T cells at 8 and 12 hours post lipofection. The control samples were: The DNA without transfection, the lipofection reagent (LTX) and EMX1 DNA. The HEK293T cells were transfected with NgAgo alone, EMX1 gDNA alone or co-transfected with NgAgo and EMX1 gDNA. A control DNA was successfully edited with CRISPR/Cas9 (+ Cas9 control) and was utilized as a negative control (- Cas9 control). The top electrophoresis gel respresents the PCR only whereas the bottom gel represents the T7E1 assay. The arrows indicate the formation of heteroduplexes for CRISPR/Cas9 genome editing. H) Western blot of NgAgo protein production with and without the co-transfection of the gDNA from 8 to 48 hours post lipofection. The staining was performed with a monoclonal Flag anti-antibody and anti-GAPDH anti-antibody. The top band represents NgAgo at 103KDa. GAPDH was utilized as a Housekeeper gene. The exposure time was 30 seconds. The controls were the lipefection agent alone (LTX), lane 1 and the gDNA EMX1 alone (lane 2). The smears under the NgAgo band show the degradation of the protein stained with the Flag tag.

**Table 1:** Generation of edited mouse blastocyst using NgAgo and CRISPR/Cas9 genome editing technologies. NgAgo circular DNA was co-injected into the mouse zygotes with various concentrations of gDNA. Cas9 Ribonucleoprotein (RNP) was injected as controls for *Sptb* targeting the same genomic sequence as NgAgo.

| Gene | Microinjection mix | Zygotes injected | Blastocysts cultured | Targeted blastocysts (%) |
| --- | --- | --- | --- | --- |
| Sptb | NgAgo 5 ng/μl + 2.5 ng/μl gDNA | 8 | 8 | 0 (0%) |
| Sptb | NgAgo 5 ng/μl + 25 ng/μl gDNA | 8 | 8 | 0 (0%) |
| Sptb | NgAgo 5 ng/μl + 50 ng/μl gDNA | 6 | 6 | 0 (0%) |
| Sptb | NgAgo 10 ng/μl + 50 ng/μl gDNA | 12 | 12 | 0 (0%) |
| Sptb | NgAgo 15 ng/μl + 50 ng/μl gDNA | 15 | 15 | 0 (0%) |
| Tet-1 | NgAgo 5 ng/μl + 25 ng/μl gDNA | 10 | 10 | 0 (0%) |
| Tet-2 | NgAgo 5 ng/μl + 2.5 ng/μl gDNA | 7 | 7 | 0 (0%) |
| Tet-2 | NgAgo 5 ng/μl + 25 ng/μl gDNA | 7 | 7 | 0 (0%) |
| Tet-3 | NgAgo 5 ng/μl + 2.5 ng/μl gDNA | 13 | 13 | 0 (0%) |
| Sptb | 50 ng/μl Cas9 + 0.6 pMol CrRNA & TracrRNA | 6 | 6 | 3(50%) |

**Table 2:**
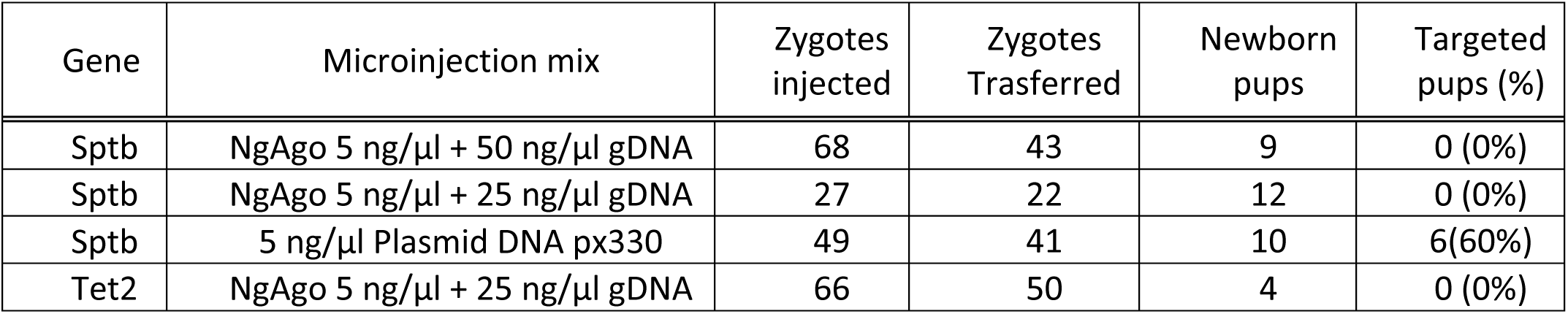
Generation of knockout mice using NgAgo and CRISPR/Cas9 genome editing technologies. Various concentrations of gDNA were co-injected with NgAgo plasmid or px330-U6-Chimeric_BB-CBh-hSpCas9 circular DNA as control.

| Gene | Microinjection mix | Zygotes injected | Zygotes Trasferred | Newborn pups | Targeted pups (%) |
| --- | --- | --- | --- | --- | --- |
| Sptb | NgAgo 5 ng/μl + 50 ng/μl gDNA | 68 | 43 | 9 | 0 (0%) |
| Sptb | NgAgo 5 ng/μl + 25 ng/μl gDNA | 27 | 22 | 12 | 0 (0%) |
| Sptb | 5 ng/μl Plasmid DNA px330 | 49 | 41 | 10 | 6(60%) |
| Tet2 | NgAgo 5 ng/μl + 25 ng/μl gDNA | 66 | 50 | 4 | 0 (0%) |

Gao et al. reported that NgAgo creates a double strand break in the DNA using a single DNA guide [2], with a reported efficiency equivalent to Cas9. Importantly, and in agreement with recently published reports [5-7], we found no evidence of the mouse and human DNA editing with NgAgo despite an efficient delivery of NgAgo and the gDNA. We did not observe an editing event in over 100 mouse embryos injected with NgAgo, giving an editing efficiency of less than 1%, contradicting the results reported in Gao et al. Interestingly, we found in the mouse embryos, mouse pups and in the HEK293T human cell line no evidence of a single indel using a T7E1 assay, and confirmed by Sanger sequencing. Qi et al reported recently a silencing role of NgAgo that may affects the phenotype of the Zebrafish embryos [6]. We have not noted such a change in phenotype in our mouse embryos. The plausible explanation for this lack of phenotype is the genes targeted were not expressed during early embryonic development [8]. Although, we did not isolate mRNA from these mouse blastocysts and we did not assess the expression of the genes.

In summary, in contradiction with Gao et al’s study, and in agreement with recently published reports, we found that NgAgo does not edit endogenous genomic DNA under physiological temperature conditions.

## Material and Methods

### Design and preparation of NgAgo, 5’-phosphorylated guide DNA (gDNA)

Four genes were targeted to design the primers and the 5’-phosphorylated oligonucleotides. These genes were the exon 26 of *Beta-Spectrin1* (*Sptb*), exon 5 of *Tet-1*, exon 3 of *Tet-2* and exon 5 of *Tet-3*. We choose gDNA previously published to be highly efficient using the CRISPR/Cas9 genome editing system. The NgAgo plasmid containing a Nuclear Localisation Signal was obtained from the Addgene repository [2]. The plasmid was cultured as per protocol and the DNA extracted using a PureLink Quick Plasmid Miniprep Kit (Invitrogen, K210010) according to the manufacturer’s instructions. 22 and 24 bp 5’ phosphorylated oligonucleotides and the amplification primers were synthetized from Integrated DNA Technologies. The sequences of these oligonucleotides are listed below (Table 3).

**Table 3:**
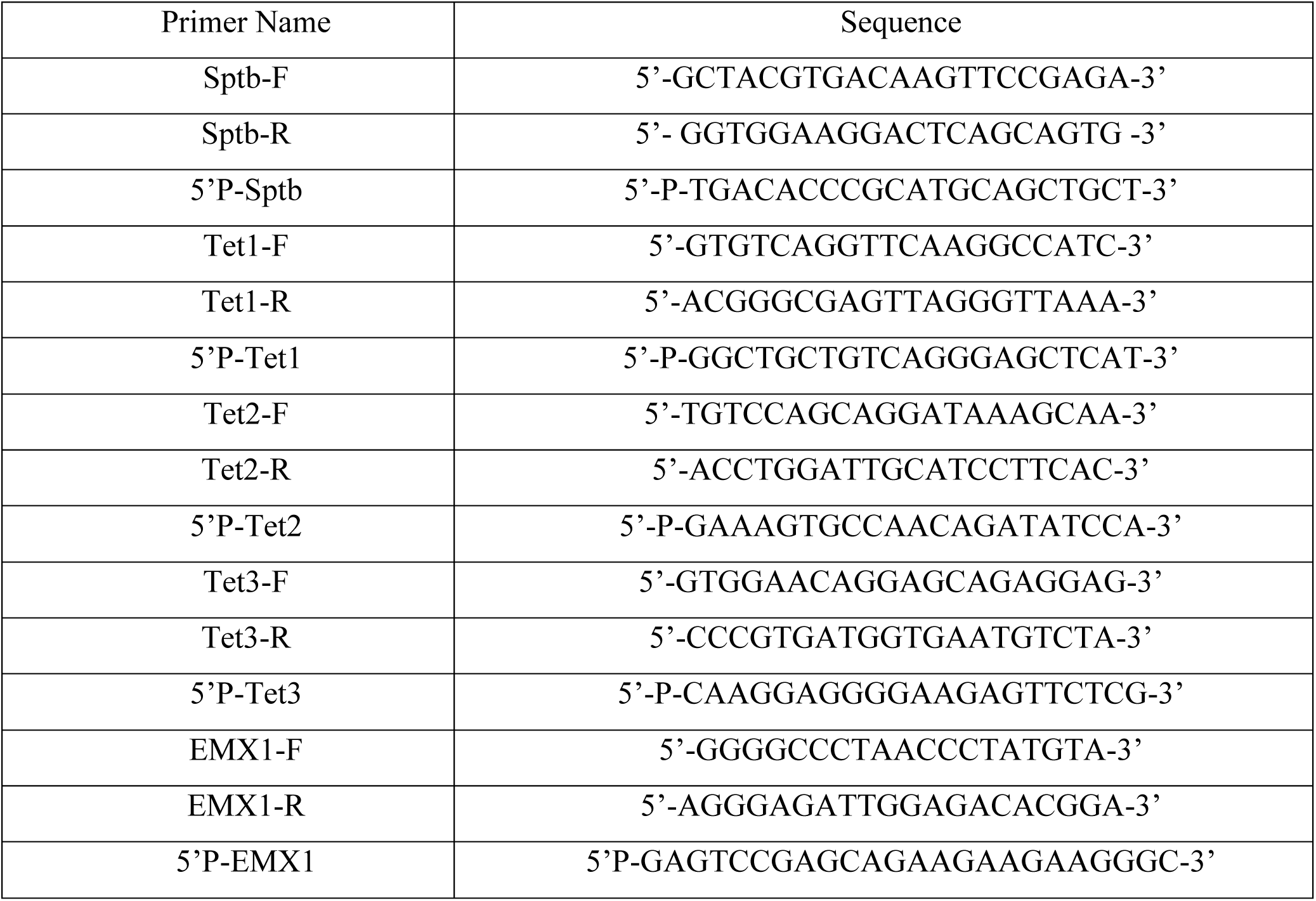
List of oligonucleotides used in this study.

### Ethics statement

All animal experiments were approved by The Australian National University Animal Experimentation Ethics Committee under the permit A2014/058 and the institutional Biosafety Committee NLRD 15.10 in accordance with the National Health and Medical Research Council (NHMRC) code of practice.

### Mouse husbandry and microinjection

C57BL/6 and recipient ICR females were purchased from Charles River laboratory and maintained in a specific pathogen-free environment at the Australian Phenomics Facility, the Australian National University. Mice were maintained on a 12h light/12h dark cycle and had ad libitum access to food and water *ad libitum*. Female C57BL/6 (3-4 weeks old, >10g) were superovulated by intraperitoneal injection of 5IU of pregnant mare serum gonadotropin (PMSG), followed by 5IU of human chorionic gonadotropin hormone (hCG) 46-48 hours later. Following injection with hCG, superovulated females were mated with stud C57BL/6 males (10-20 weeks old). The embryos were collected from oviducts approximately 45 hours after the last injection and held in M16 medium (Sigma M7292) overlaid with mineral oil at 37°C and 5% CO_2_ until injection.

Microinjection was performed in M2 medium (Sigma, M7167) under mineral oil using an inverted microscope (Leica DMi8) and micromanipulators. Pronuclear injection of fertilized zygotes was performed with the following mixes: circular plasmid DNA at 5 ng/μl and 5’-P oligonucleotide at 2.5, 25 and 50 ng/μl. For Sptb CRISPR injection, the gRNA was cloned into px330 vector [9] obtained from Addgene (ID 42330) using the following oligonucleotides 5’- CACCGTGACATGTGGGCGGACCTGC - 3’ and 5’ – AAACGCAGGTCCCCACATGTCAC – 3’. 5 ng/μl of px330 circular plasmid was injected into the fertilized zygotes by pronuclear injection to obtained live mice. For blastocysts culture, fertilized zygotes were microinjected using the following mix: 50 ng/μl of Cas9 purified protein from PNA BIO (Thoussand Oaks, CA), 0.6 pMol of CrRNA 5’ – UGACAUGUGGGCGGACCUGGUUUAGAGCUAUGCUGUUUUG - 3’ and 0.6 pMol of TracRNA. The CrRNA and TracRNA were complexed with the Cas9 protein by incubating at 37°C for 10 minutes. Microinjected zygotes were cultured overnight in M16. Resulting two-cell embryos were surgically transferred into the ampulla of pseudo-plugged ICR female recipients (8-12 weeks old) or cultured in M16 media for 4 days at 37°C.

### Genotyping

A subset of the microinjected zygotes was cultured for 4 days to the blastocyst stage in M16 medium overlaid with mineral oil at 37°C and 5% CO_2_. The other zygotes were cultured for 24 hours and surgically transferred into the surrogate mouse ampulla. The mice were maintained and the resulting pups were maintained and genotyped 15 days after birth. DNA was extracted from the blastocysts at day 5 or live mouse pups over 15 days old using a crude DNA extraction protocol. In short, the blastocysts were lysed in Tris-EDTA-Tween lysis buffer (50mMTris HCl, pH8.0, 0.125mM EDTA, 2% Tween 20) with 1μl of proteinase K (20 mg/ml in 10mM Tris chlorate, 0.1 mM ethylenediaminetretaacetic acid (EDTA) pH 8.0) and incubated at 56°C for an hour before being denatured at 95°C for 10 minutes. We amplified regions encompassing the gDNA with 2x MyTaq HS mix (Bioline, cat no. BIO-25045) under the following PCR conditions: 95°C for 3 minutes followed by 35 cycles (95°C for 15”, 58°C for 15” and 72°C for 20”) and 72°C for 3 minute. The PCR products were checked on a 1.5% electrophoresis gel. The PCR products were purified with ExoSAP-IT® (affymetrix, Cat no. 78202), or cut from the gel and purified using the Wizard® SV Gel and PCR Clean-Up System (Promega, Cat no. A9282) kit according to the manufacturer’s instructions. The Sanger sequencing was conducted at the Biomedical Resource Facility at the John Curtin School for Medical Research, The Australian National University.

### T7 endonuclease assay

After PCR amplification, the PCR fragments were hybridized and digested with a T7 endonuclease (NEB, Cat no. M0302S) for 15-30 minutes at 37°C. After digestion, the enzymatic reaction was stopped using 1 μl of 0.25 M EDTA and the digested product run on a 1.5% agarose gel alongside the undigested PCR product as a control.

### Construction of a Flag-nls-NgAgo-GK plasmid

The nls-NgAgo-GK plasmid was a gift from Chunyu Han (Addgene plasmid 78253). nls-NgAgo-GK plasmid was digested overnight with AleI (NEB, cat no. R0634S). Following the digestion a forward (5’- TGGACTATAAGGACCACGACGGAGACTACAAGGATCATGATATTGATTACAAAGACGATGACGATAAGA-3’) and a reverse (5’ - TCTTATCGTCATCGTCTTTGTAATCAATATCATGATCCTTGTAGTCTCCGTCGTGGTCCTTATAGTCCA-3’) oligonucleotide sequence encoding for a flag tag were annealed and ligated into the nls-NgAgo-GK plasmid upstream to the SV40 Nuclear Localization Signal (nls). Sanger sequencing was used to assess the correct integration of the flag-tag in frame with the start codon. The plasmid was transformed into heat-chock competent BL21 *E. coli*. The plasmid is being deposited at Addgene (Plasmid #73681) and will be available to the community.

### Cell culture and transfection

HEK293T cells were obtained from ATCC (CRL-11268). The cells were maintained in Dulbecco’s Modified Eagle Medium (DMEM) (Sigma, Cat. No. D6546) supplemented with 10% heat-inactivated fetal bovine serum (Sigma, Cat. No. 12003C), 2mM L-Glutamine/1%Penicilin/Streptomycin solution (Thermofisher, Cat. No. 10378016) and incubated at 37°C with 5% CO_2_. The cells were seeded at 2x10^5^ cells per well in 2.5 mL of medium in a 6-well plate, to reach 60-70% confluency immediately prior to transfection. Per sample well, 3.0ug of plasmid DNA (Flag-nls-NgAgo-GK) and/or 0.5μg of 5’ phosphorylated EMX1 guide oligo was added to 150uL of basal DMEM (without additives) with 3.5uL Plus Reagent (Invitrogen, Thermofisher, Cat. No. 15338100). 150uL of this diluted DNA mixture was added to 150uL of DMEM (without additives) and 12uL of Lipofectamine LTX® Reagent (Invitrogen, Thermofisher, Cat. No. 15338100). The mixture was incubated for 15 minutes at room temperature to form DNA-Lipofectamine LTX® Reagent complexes. After incubation, 250uL of the DNA-Lipofectamine complex was added drop-wise to each well, and the plate was gently rocked. Negative controls constituted a ‘cell growth’ control, with no DNA or lipofectamine reagents, and a ‘lipofectamine only’ control with no plasmid or gDNA. Transfected cells were incubated at 37°C in a 5% CO_2_ incubator for 8, 12, 24 and 48 hours post-transfection before collecting proteins to assay for transgene expression, and extracting genomic DNA for genotyping assays.

### Genomic DNA extraction and protein extraction from HEK293T cells

The genomic DNA was isolated and purified using ISOLATE II Genomic DNA Kit (Bioline, BIO-52066) according to the manufacturer’s instructions. To extract proteins from HEK293T cells, the cells were incubated with RIPA buffer (5M NaCl, 0.5M EDTA, pH 8.0, 1M Tris, pH8.0, 1% TritonX-100, 10% sodium deoxycholate, 10% SDS, 1x Protease Phosphatase Inhibitor) with gentle rocking on ice for 30 minutes. The cells were scraped off the plates to dislodge lysate. Lysate was centrifuged at 13,000g for 5 minutes at 4°C. The supernatant of lysate was stored at -20°C.

### Western blotting

The nuclear and cytoplasm lysate samples were denatured at 95°C for 5minutes in 1x Laemmli buffer prior to loading onto a 4-15% SDS-PAGE (4–15% Mini-PROTEAN® TGX^TM^ Precast Protein Gels, Bio-Rad #4561085). 10uL Precision Plus Protein^TM^ Kaleidoscope^TM^ Prestained Protein Standards (Bio-Rad, #1610375) or 10-15uL of samples were loaded on to the gel. The gel was eletrophorised for 35-40mins at 200mV, in 1x running buffer (25mM Tris-Base, 190mM glycine, 0.1% SDS, pH 8.3). Proteins were transferred onto a nitrocellulose membrane (Bio-Rad, Cat no. 162- 0115) at 400mA, 300V for 1 hour and 30 mins at 4°C, in 1x transfer buffer (25mM Tris-Base, 190mM glycine, 20% methanol, pH 8.3). Primary antibodies (anti-FLAG 1:1000, Sigma Cat no. F1804-200UG, or anti-GAPDH1:1000, Millipore Cat no MAB374) were diluted in 1% skim milk in PBS and incubated for 1 hour at RT with shaking. After incubation, membranes were washed with 1xPBS+0.1%Tween for three times 5 mins and once for 10 mins. After washing, membranes were incubated with secondary antibody (1:5000 goat anti-mouse Ig-HRP, Sigma Cat no. A44161ML) diluted in 1% skim milk in PBS, for 1 hour at RT with shaking, and then washed again as above. Membrane was visualised under chemiluminescence with HRP substrate (Millipore, Cat no. WBLUF0500) at various exposure times.

### Analysis and Statistics

Using ImageJ 1.05i software, Western blot membranes were analyzed for the mean band intensity for anti-FLAG (corresponding to the expression of NgAgo), as well as for the loading control anti-GAPDH. The relative abundance of anti-FLAG was determined as a ratio of the loading control to produce a FLAG:GAPDH ratio. These ratios were then normalized to the negative control, containing no NgAgo. These values were then analysed as listed below.

A Two-Way ANOVA using Tukey’s multiple comparisons test was performed to quantify changes in protein expression over time for each treatment. Paired T-tests were performed to compare protein expression over time between NgAgo samples with and without gDNA. A two-tailed Wilcoxon test was performed to examine changes in degradation between samples with or without gDNA. All analyses were conducted using the GraphPad software, Prism 7, with significance at P≤0.05.

## Acknowledgments

This work was supported from the National Collaborative Research Infrastructure (NCRIS) through the Australian Phenomics Network in Australia. The funders have no role in study design, data collection and analysis, decision to publish or preparation of the manuscript. We would like to thank all the comments, anonymous or not on the initial blogpost, on Twitter and/or on a Google discussion group dedicated to genome editing technology that have led to stimulating, interesting and fruitful discussions on this technique. Finally we would like to thanks 3 anonymous reviewers for their helpful comments that has improved the quality of the manuscript.

## Author contribution

Designed the experiments: NCK, JL, LJ and GB. Performed the experiments NCK, JL, LJ. Analyzed the data: NCK, JL, LJ and GB. Wrote the manuscript GB with the input from NCK, JL and LJ. All the authors read and approved the manuscript.

---

**Supplementary Figure 1.**
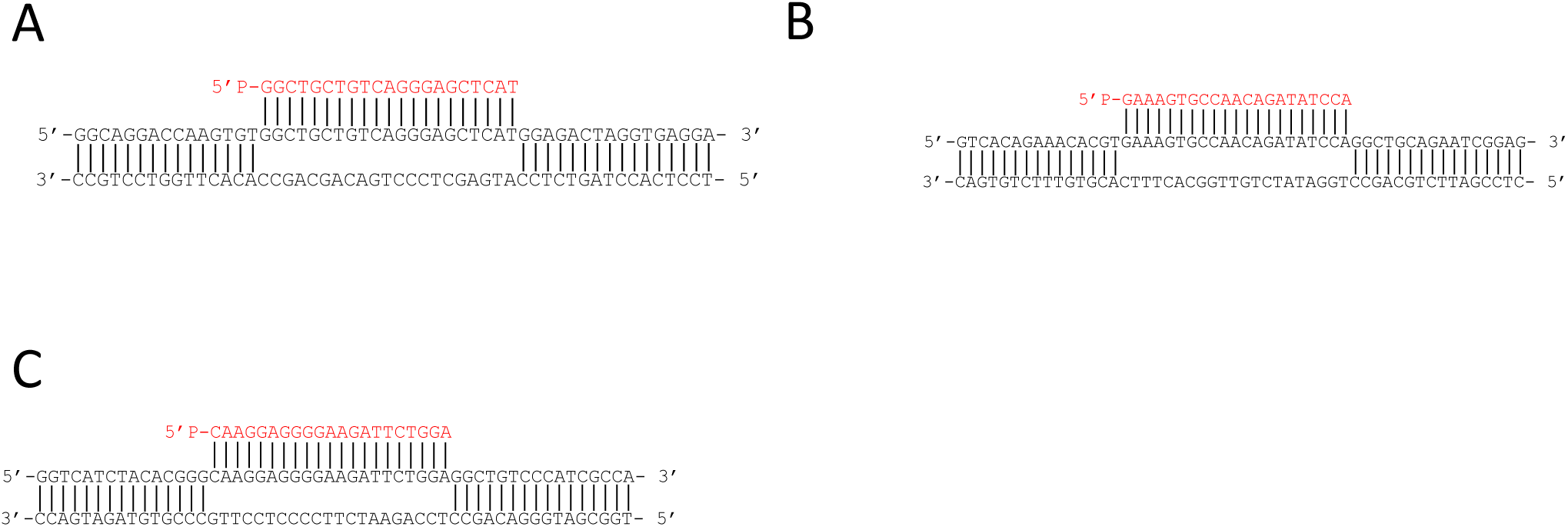
Khin et al.

**Supplementary Figure 2.**
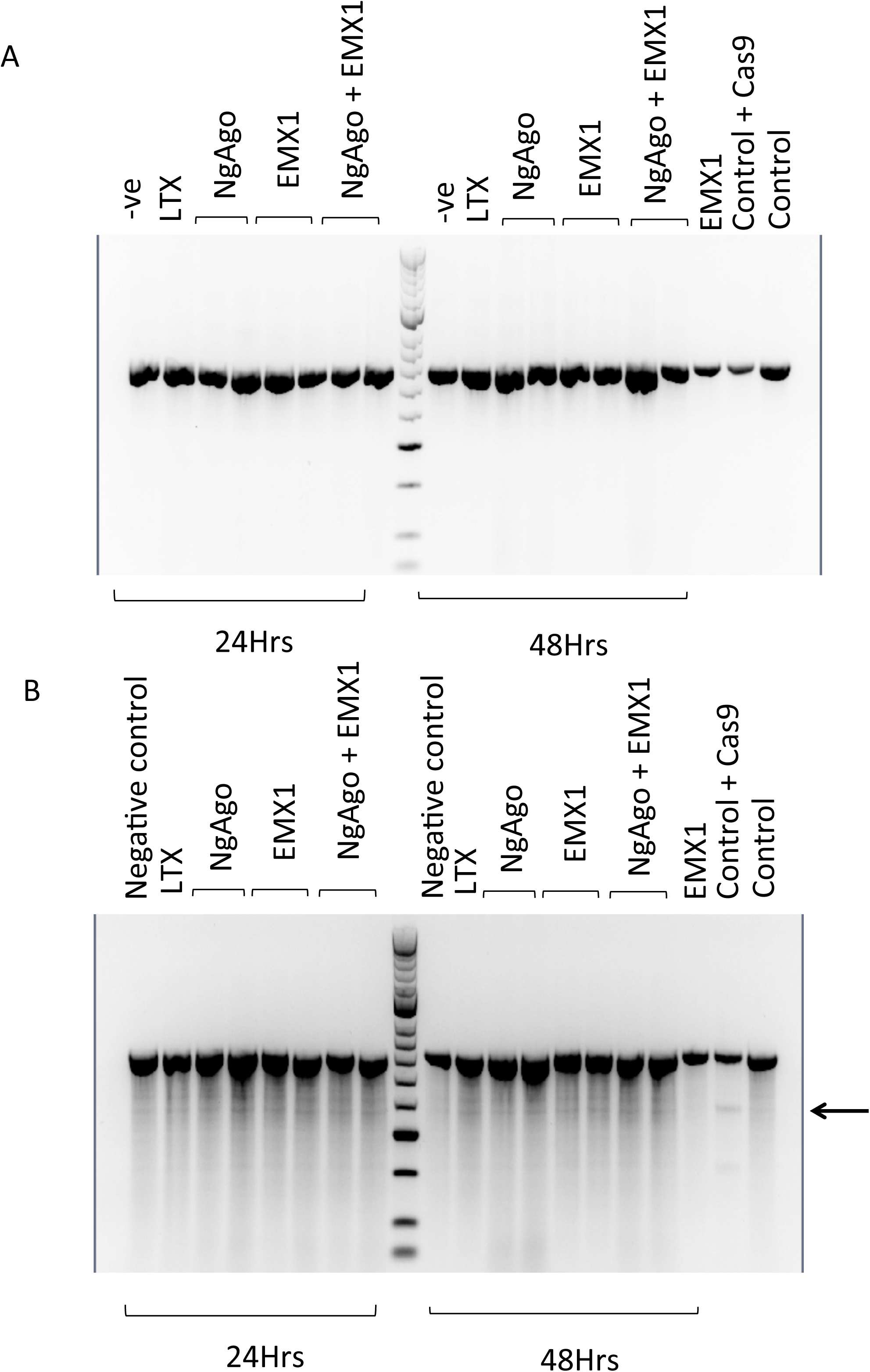
Khin et al.

**Supplementary Figure 3.**
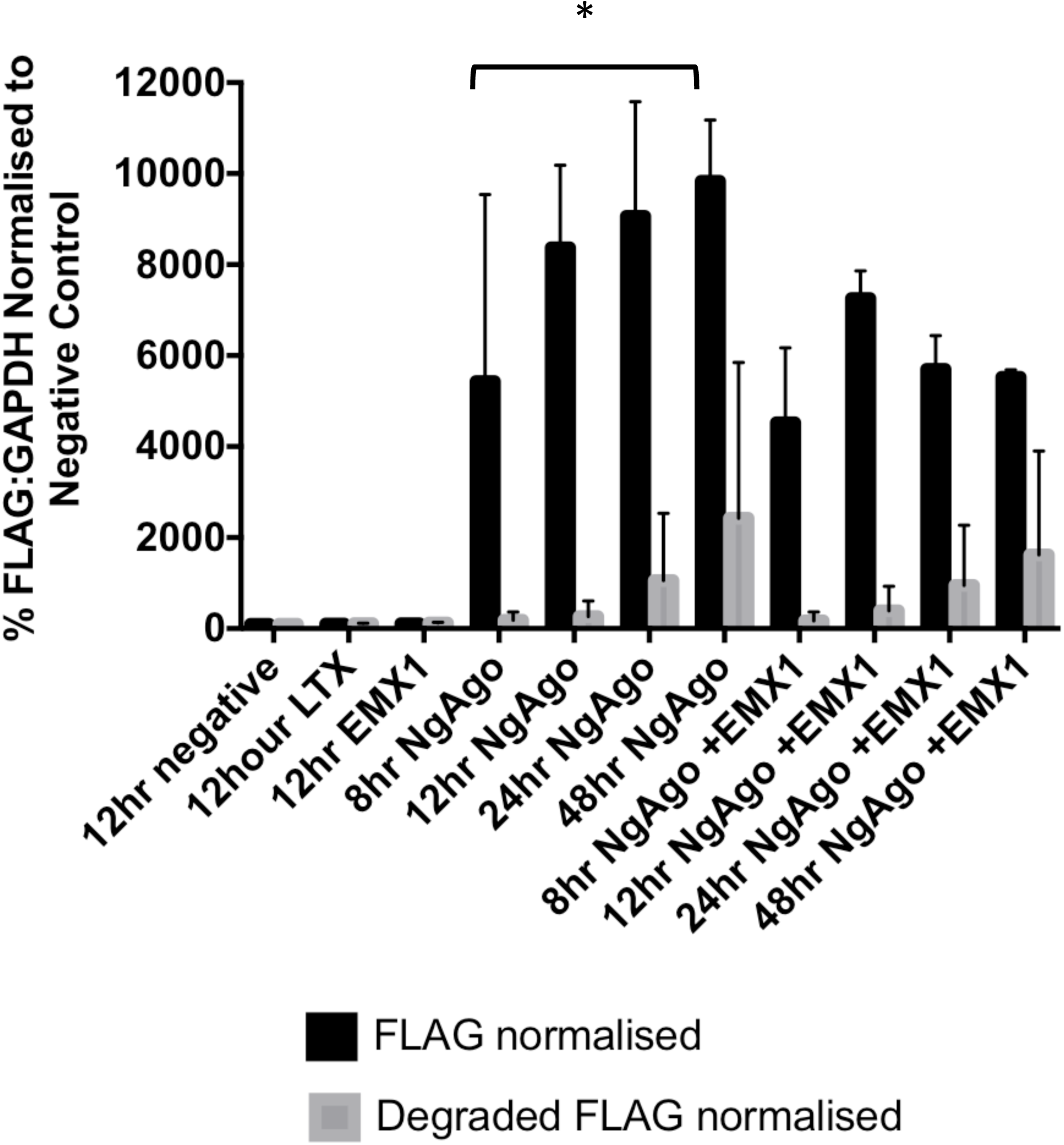
Khin et al.

